# Single Cell Pluripotency Regulatory Networks

**DOI:** 10.1101/062422

**Authors:** Patrick S. Stumpf, Rob Ewing, Ben D. MacArthur

## Abstract

Embryonic stem (ES) cells represent a popular model system for investigating development, tissue regeneration and repair. Although much is known about the molecular mechanisms that regulate the balance between self-renewal and lineage commitment in ES cells, the spatiotemporal integration of responsive signalling pathways with core transcriptional regulatory networks are complex and only partially understood. Moreover, measurements made on populations of cells reveal only average properties of the underlying regulatory networks, obscuring their fine detail. Here, we discuss the reconstruction of regulatory networks in individual cells using novel single cell transcriptomics and proteomics, in order to expand our understanding of the molecular basis of pluripotency, including the role of cell-cell variability within ES cell populations, and ways in which networks may be controlled in order to reliably manipulate cell behaviour.

## 1 Introduction

Our understanding of how pluripotency has grown tremendously since pioneering studies described the derivation and *in vitro* culture of pluripotent stem cells (PSCs) [1–8]. It is now well known that PSCs display two characteristic features: 1) indefinite self-renewal in vitro, and 2) tri-lineage commitment to ectoderm, endoderm and mesoderm, once released from the self-renewing regime. Knowledge of both these properties has been predominantly generated from aggregates of cellular material, and therefore represents the average behaviour of hundreds or thousands of cells. Nevertheless, from a wealth of regulatory relationships, a limited set of core transcription factors have been inferred and validated, resulting in the construction of a now reliable regulatory model for the pluripotent state [9]. However, more recent measurements at the single cell level have highlighted the presence of significant biological variability and heterogeneity among clonal PSC populations [10], suggesting that subtle cellcell variations in network configurations may have an important role in regulating pluripotency [11–15]. These results stress the importance of reconstructing regulatory networks at the individual cell level in order to uncover the refined mechanisms that balance self-renewal and differentiation *in vitro* and cellular propensities for different developmental states. In this article, we sketch out our current understanding of the integrated regulatory network (IRN) that controls the transient developmental state of pluripotency and discuss the ways in which more refined single cell regulatory networks are enhancing our understanding of PSC states and the ways in which PSCs balance self-renewal and lineage commitment at the individual cell level.

## 2 Combinatorial control of pluripotency by regulatory networks

The pluripotent state in mouse and human cells is regulated by a number of integrated regulatory networks (previously reviewed [16–20]), including transcriptional [21], epigenetic [22], signalling [23] and metabolic [24] sub-networks (represented schematically in Figure 1). In the presence of defined extrinsic stimuli [25,26], the pluripotent state is maintained by a cell-intrinsic set of transcription factors (TFs) that constitute a self-sustaining gene regulatory network (GRN) that is rich in feedback [27]. Central to this GRN lies a core network of TFs composed of Oct4, Sox2, Nanog, with significant support of secondary factors such as also Klf4, Myc and Lin28 [28]. Combinations of these TFs were originally found to revert the cell identity of terminally differentiated somatic cells towards the pluripotent cell identity [6–8], however subsets of these factors are also sufficient to reconstitute pluripotency in somatic cells [29,30]. The members of this core GRN interact with a range of auxiliary transcription factors [31–36], which collectively control transcription of a large number of genes either directly, by binding to gene promoters [21,33,37,38], or indirectly, by mediating the effects of epigenetic remodelling complexes [20,39,40], which help maintain pluripotency by producing a permissive chromatin state that allows for widespread nonspecific transcription [41], in which important developmental genes are sporadically expressed at low levels, yet remain poised for robust expression under the appropriate differentiation cues [42–44]. To buffer this “noisy” environment, a network of microRNAs [45–48] and ribosome specific mechanisms [49], ensure appropriate protein levels are robustly maintained. In addition to these cell-intrinsic regulatory mechanisms, a layer of signalling pathways integrates cell-extrinsic information to the central pluripotency GRN. While the core transcriptional circuitry is broadly similar in mouse and human cells [50], mouse embryonic stem cells (mESCs), mouse epiblast stem cells (mEpiSCs) and human embryonic stem cells (hESCs) display marked differences in their dependence on extrinsic signalling factors. In mESCs, Lif/Stat signalling [51,52], Bmp [53] and canonical Wnt [25] promote self-renewal, while Fgf/Erk signalling disrupts pluripotency [25,54–56]. In contrast, hESCs and mEpiSCs require Activin and Fgf [57,58] signalling for self-renewal and cells in this “primed” pluripotent state undergo differentiation when exposed to Bmp [58], while Lif/Stat signalling has no measureable effect on their self-renewal *in vitro* [59]. Importantly, the flow of information between signalling and transcriptional regulatory networks is not one-way: signalling networks mediate external environmental information to the core GRN, while the core GRN affects the expression of the pathway components themselves, or of key miRNAs that in turn regulate signalling pathway components [60]. Collectively, these reports indicate that pluripotency is regulated by mechanisms that act at both the transcriptional and translational levels and involve layers of combinatorial regulatory control, including complex feedback relationships between transcriptional, epigenetic and signalling mechanisms. However, while this model has been tremendously successful, much of this information has been inferred from bulk properties of large ensembles cells. Within individual cells, regulatory networks may adopt a variety of different states and may deviate dramatically from this ensemble model. Thus, a better understanding of how cell-cell variation in network structure affects cell population function is now needed.

**Figure 1.**
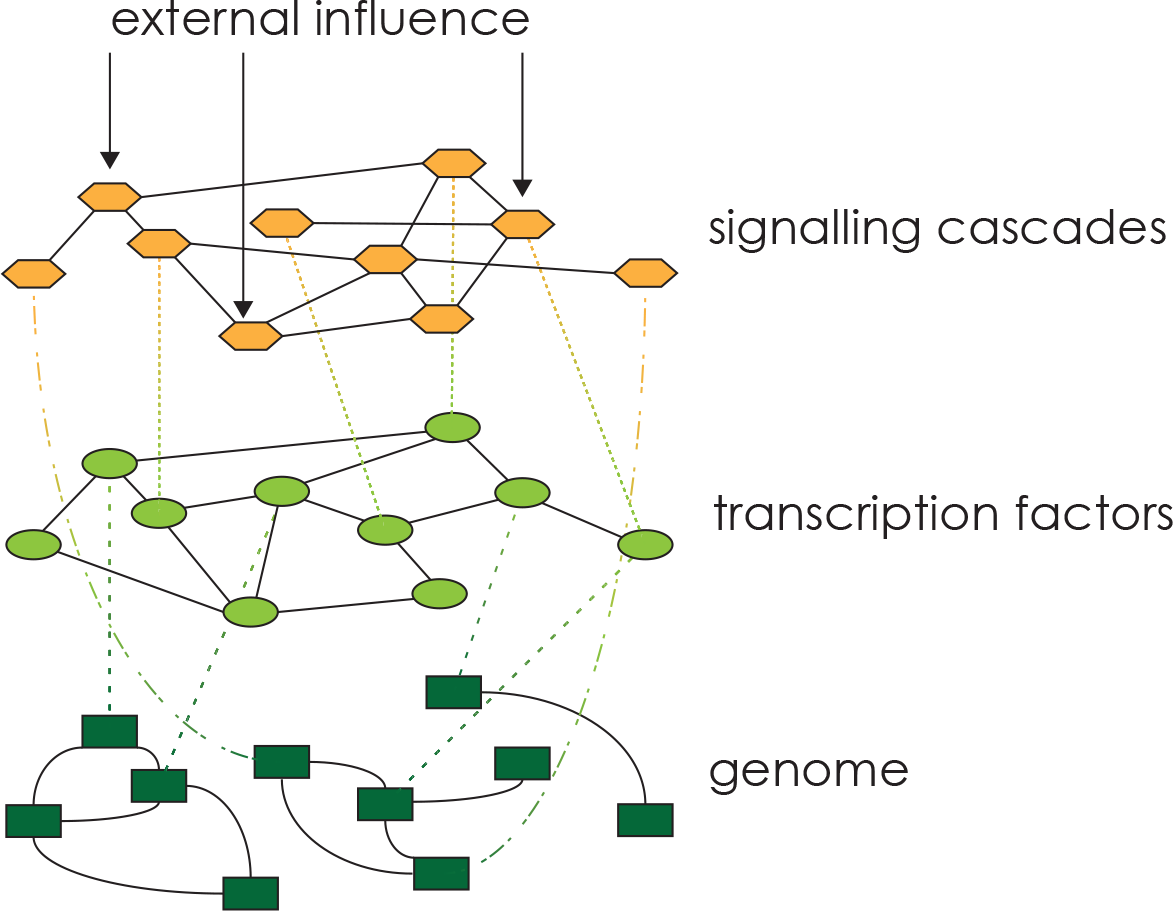
Pluripotency is regulated by a multifaceted network composed of several sub-networks, such as signalling cascades (yellow), interactions among transcription factors (light green), or epigenetic networks (dark green) that are characterised by qualitatively different interactions. However, each subnetwork is connected, for instance through the directional flow of information from genome to the proteome, but also via complex feedback interactions between the different “layers”. Collectively, these sub-networks form an integrated regulatory network (IRN).

## 3 Regulatory networks at the single cell level

In contrast to ensemble networks from bulk cell material, single cell measurements of co-expression patterns are able to reveal a more nuanced picture of regulatory networks within cell populations [61]. Traditionally, flow-cytometry (FC) has been used quantify co-expression patterns of individual cells. However, FC methods are intrinsically limited in the number of factors that can be co-assessed (currently up to about 18), mainly due the spectral resolution of fluorescently labelled antibodies [62]. These limitations are gradually being overcome, for instance using new methods such as Cytometry by Time of Flight (CyTOF), which, at present, is able to quantify co-expression of up to approximately 50 different proteins in individual cells via immuno-labelling with elemental isotopes [62–64]. Similarly, complementary nucleic-acid-based techniques such as RNA-FISH [65,66], qRT-PCR [67] and RNA-seq [68] are also able to assess multi-dimensional transcript expression patterns at single cell resolution. These emerging single cell technologies are now enabling broad expression profiles across a large number of cells to be obtained [69], from which single cell regulatory networks can be inferred [70]. Table 1 summarizes some notable uses of these methods to profile PSCs.

**Table 1.**
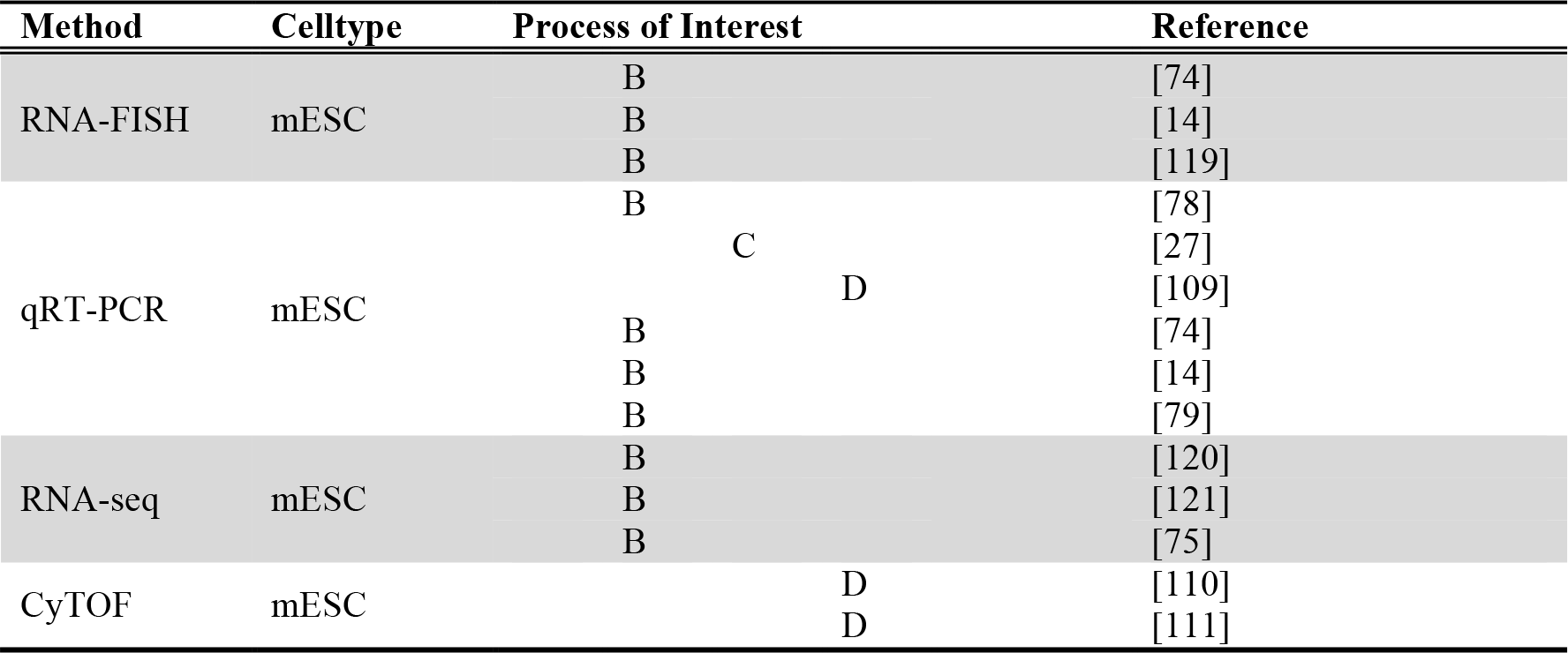
Available datasets for reconstructing single cell regulatory networks. Cellular processes of interest are: **A** developmental origins of pluripotency in the inner cell mass; **B** alternative pluripotent states; **C** exit from pluripotency and; **D** establishing pluripotency ectopically in somatic cells.

Using conventional methods such as FC, evidence of heterogeneity in clonal cell populations has accumulated for individual factors of the integrated pluripotency regulatory network [14,71–75] and it is generally agreed that variable quantities of mRNA and protein can be present within individual cells of a clonal cell population, for instance due to noise inherent to transcription and translation [76,77]. For important members of the IRN such variation can strongly affect the efficiency with which cell-extrinsic stimuli are processed by individual cells and thereby drive widespread differences in multivariate expression patterns within clonal populations (for example amongst the descendants of individual PSCs) [15]. Moreover, variability in expression of central regulatory factors can also affect network structure by differentially activating particular sub-networks of the IRN within individual cells [78]. Such structural changes have been shown to have an important role in regulating cell-cell variability in PSC fate changes, for example upon Nanog withdrawal [27,79]. Ultimately, differential expression of important sub-networks may impose constraints on cellular signal processing, which in turn restrict the degrees of freedom of cell-fate decisions. Although yet fully demonstrated, it is likely that similar mechanisms, involving subtle variations in network structure between in individual cells, are important for regulating the collective dynamics of PSC populations [80].

## 4 Plasticity of pluripotency networks in development and reprogramming

Regulatory networks controlling a particular cellular identity undergo dramatic changes as cells progress from one developmental state to another [81,82]. Such changes can be exploited to classify cellular identities based on properties of their underlying regulatory network [83,84]. Four instances of cellular identity changes are of particular interest with respect to understanding cell-cell variability and network-plasticity in pluripotency: three of these instances are associated with the native developmental programmes, starting with blastulation (the origin of pluripotency) and proceeding through gastrulation (establishment of different pluripotent states, followed by exit from pluripotency); while the fourth is related to the reverse process of establishing the pluripotent regulatory network in somatic cells during cellular reprogramming. In all four cases, variation in cell-cell expression of regulatory networks has an important role.

### A *Network plasticity during blastulation*

During pre-implantation development, the inner cell mass (ICM) forms two strata, the epiblast (EPI) and primitive endoderm (PE), in preparation to the formation of the embryo proper from the EPI. Initially, mosaic expression patterns of central regulatory factors for EPI (Nanog) and PE (Gata6) emerge seemingly at random [85]. The apparent spatial randomness of this process suggests that cell intrinsic stochastic mechanisms are responsible for the initial EPI-PE stratification process [86], while sub-networks centred on Nanog or Gata6 subsequently reinforce these initial stochastic variations before cell re-arrangements, coordinated through juxtacrine signalling, lead to tissue-organisation into the two strata [87,88]. However, other evidence suggests that the mosaic expression of Nanog and Gata6 is preceded by asymmetric cell division leading to an unequal distribution of Fgf-signalling components [89–91]. The reconstruction of single-cell regulatory networks could be instrumental in consolidating both models by inferring the logical sequence of events from unbiased single cell expression data.

### B *Network plasticity during the naive pluripotent to primed pluripotent transition*

Two pluripotent states exist that display distinct differences in their IRN [92]: a naïve pluripotent state present in the EPI of the pre-implantation embryo, from which mESCs are derived; and a primed state, characteristic of the late stage of the EPI in the post-implantation embryo (the egg cylinder in mice), from which mEpiSCs are derived [26]. While the transition from the naïve state to the primed state corresponds to the natural developmental progression in the embryo, the primed states can be artificially reverted to the naïve state *in vitro* only through ectopic expression of Klf4 in mEpiSCs [93], or Nanog and Klf2 in hPSCs [94]. These observations reveal a remarkable property: only few key nodes are necessary to alter the processing logic of the IRN towards accepting contrasting extrinsic signalling inputs (i.e. LIF and BMP in mESCs versus Activin and Fgf/Erk in mEpiSCs) in order to arrive at the same outcome: self-renewal. The precise rewiring of signalling pathways into the GRN, characteristic for the alternative pluripotent states, may be inferred from single-cell expression data.

### C *Decay of the IRN: From pluripotency and lineage commitment*

In development, the transient pluripotent state ceases with the formation of the germ layers during gastrulation. The spatial organisation of the peri-implantation embryo contains various localised sources of signalling molecules [95] and it has been demonstrated that such localised extrinsic signals can cause asymmetric division, leading to two daughter cells with different sets of active signalling networks and GRN components [96]. If key GRN components such as Nanog are lost – with accompanying changes in transcription factor binding [97] and chromatin reorganisation [98] – then the self-sustaining properties of core GRN in one daughter may be compromised, leading to destabilisation of the pluripotent state and spontaneous differentiation [27]. Thus, changes in expression of key factors subsequent to cell division can lead to divergent fates in paired daughter cells, via reorganization of intracellular regulatory networks (see Figure 2).

**Figure 2.**
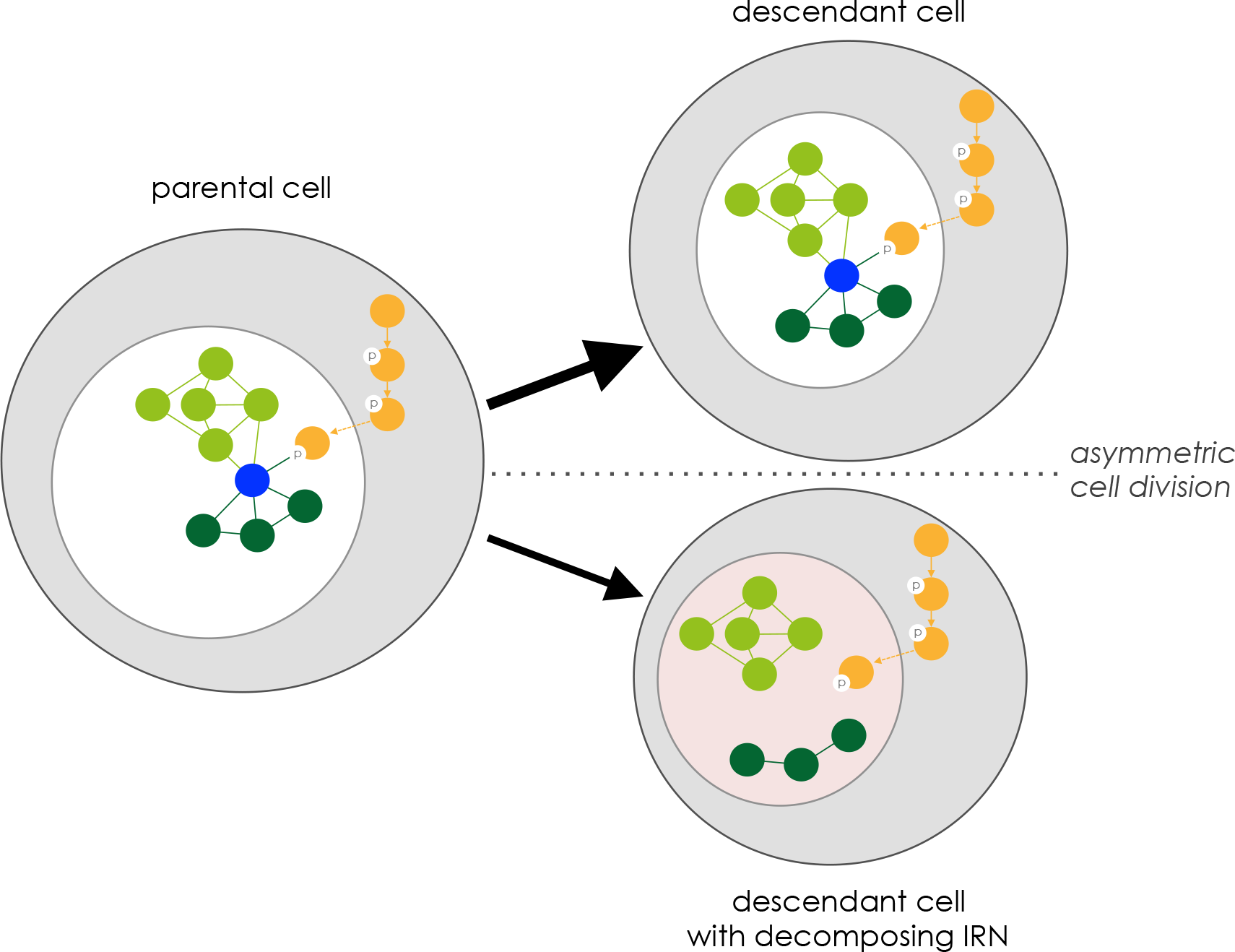
Divergence of sister cell fates upon an asymmetric division. Differential inheritance of factors associated with important nodes in the IRN can lead to profound differences in regulatory network structure in the daughter cells. For instance, loss of a central transcription factor (blue) in one of the daughter cells can lead to the dissociation of signalling cascades (yellow), transcriptional networks (light green) and epigentic networks (dark green).

### D *Induced pluripotency*

The most dramatic changes to the IRN occur during reprogramming *in vitro* [99]. The principle of cellular reprogramming is to establish pluripotency in somatic cells by transiently inducing the activity of key parts of the self-sustaining pluripotency network, for instance by ectopic overexpression of core factors in direct reprogramming [6–8]. The more components a somatic cell IRN and the pluripotent IRN have in common, the fewer factors are required for this identity-remodelling [29,30]. Moreover, redundancies in the pluripotency IRN allow the replacement of individual factors without affecting the final cell identity [100]. Full reprogramming typically takes a number of weeks, and along the way to the pluripotent state, cells transition through a range of intermediate signalling [101] and chromatin states [102,103]. Although the specific trajectory taken depends upon the cells initial identity, it has been shown, using a drug inducible system for comprehensive dedifferentiation [104], that the progeny of the majority of cell types are able to undergo an identity conversion towards pluripotency [105]. Although early reports indicated that reprogramming is a stochastic process [105–107], in the case that overexpression of key factors in supplemented with additional knockdown of Mbd3 deterministic reprogramming with synchronised emergence of pluripotent colonies has been observed [108], suggesting that reprogramming progresses through a fixed sequence of remodelling events. Thus, the balance between stochastic and deterministic mechanisms has yet to be fully elucidated. Single cell based expression data, addressing the sequence of reprogramming events have very recently become available [109–111]. Using these data and similar experiments to study the corresponding topological changes to the IRN that occur during this transition, will likely inform important questions surrounding the nature of reprogramming.

## 5 Controllability of single cell networks

In summary, many efforts have been made to decipher the components within the IRN that control average cell behaviour [112–114]. The structure of these ensemble networks can explain this population-level behaviour [9], however, due to variability in the expression levels, and, necessarily, cell-to-cell differences in the IRN, the response to the provided stimuli will vary greatly among individual cells, leading, for instance, to impurities in the resulting cell population following ‘directed’ differentiation, or, as another example, incomplete cellular reprogramming. An intuitive strategy to develop better experimental protocols is to identify important driver nodes, in order to reduce the undesirable by-products that emerge from these processes. These driver nodes are the set of nodes that must be manipulated in order to control a system completely [115], for instance to steer the IRN of a particular cell state into the desired alternative state. In the absence of a full understanding of the structure and dynamics of the IRN this is a challenging task, although recent developments in the theory of controllability of networks may help [116]. One strategy to address this problem is the computational inference of regulatory networks from single cell data. For this purpose, a number of methods are available [117,118]. Individual studies that have already started to adopt similar strategies employed Boolean networks to reconstruct pluripotent IRN from single cell qPCR data [79] or Bayesian network inference in order to extract the sequence of events during reprogramming [109]. We predict that such methods – which combine high-throughput single cell profiling, with advanced network analysis routines – will lead to a more complete understanding of pluripotency in general and the development of better protocols for stem cell maintenance, differentiation and reprogramming in particular.

## Conflict of Interest Statement

The authors have declared no conflict of interest.

## References

[1] Martin, G.R., Isolation of a pluripotent cell line from early mouse embryos cultured in medium conditioned by teratocarcinoma stem cells. PNAS 1981, 78, 7634–7638.

[2] Evans, M.J., Kaufman, M.H., Establishment in culture of pluripotential cells from mouse embryos. Nature 1981, 292, 154–156.

[3] Thomson, J.A., Itskovitz-Eldor, J., Shapiro, S. S., Waknitz, M. A., et al., Embryonic stem cell lines derived from human blastocysts. Science 1998, 282, 1145–1147.

[4] Tesar, P. J., Chenoweth, J. G., Brook, F. A., Davies, T. J., et al., New cell lines from mouse epiblast share defining features with human embryonic stem cells. Nature 2007, 448, 196–199.

[5] Brons, I.G.M., Smithers, L. E., Trotter, M.W.B., Rugg-Gunn, P., et al., Derivation of pluripotent epiblast stem cells from mammalian embryos. Nature 2007, 448, 191–195.

[6] Takahashi, K., Yamanaka, S., Induction of pluripotent stem cells from mouse embryonic and adult fibroblast cultures by defined factors. Cell 2006, 126, 663–676.

[7] Takahashi, K., Tanabe, K., Ohnuki, M., Narita, M., et al., Induction of pluripotent stem cells from adult human fibroblasts by defined factors. Cell 2007, 131, 861–872.

[8] Yu, J., Vodyanik, M. A., Smuga-Otto, K., Antosiewicz-Bourget, J., et al., Induced pluripotent stem cell lines derived from human somatic cells. Science 2007, 318, 1917–1920.

[9] Dunn, S.-J., Martello, G., Yordanov, B., Emmott, S., Smith, A. G., Defining an essential transcription factor program for naïve pluripotency. Science 2014, 344, 1156–1160.

[10] Huang, S., Non-genetic heterogeneity of cells in development: more than just noise. Development 2009, 136, 3853–3862.

[11] MacArthur, B. D., Lemischka, I. R., Statistical mechanics of pluripotency. Cell 2013, 154, 484–489.

[12] Torres-Padilla, M.-E., Chambers, I., Transcription factor heterogeneity in pluripotent stem cells: a stochastic advantage. Development 2014, 141, 2173–2181.

[13] Abranches, E., Guedes, A.M.V., Moravec, M., Maamar, H., et al., Stochastic NANOG fluctuations allow mouse embryonic stem cells to explore pluripotency. Development 2014, 141, 2770–2779.

[14] Kumar, R. M.,Cahan, P., Shalek, A. K., Satija, R., et al., Deconstructing transcriptional heterogeneity in pluripotent stem cells. Nature 2014, 516, 56–61.

[15] Filipczyk, A., Marr, C., Hastreiter, S., Feigelman, J., et al., Network plasticity of pluripotency transcription factors in embryonic stem cells. Nat Cell Biol 2015, 17, 1235–1246.

[16] Jaenisch, R., Young, R., Stem cells, the molecular circuitry of pluripotency and nuclear reprogramming. Cell 2008, 132, 567–582.

[17] MacArthur, B. D., Ma'ayan, A., Lemischka, I. R., Systems biology of stem cell fate and cellular reprogramming. Nat Rev Mol Cell Biol 2009, 10, 672–681.

[18] Pera, M. F., Tam, P.P.L., Extrinsic regulation of pluripotent stem cells. Nature 2010, 465, 713–720.

[19] Young, R. A., Control of the embryonic stem cell state. Cell 2011, 144, 940–954.

[20] Orkin, S. H., Hochedlinger, K., Chromatin connections to pluripotency and cellular reprogramming. Cell 2011, 145, 835–850.

[21] Boyer, L. A., Lee, T. I., Cole, M. F., Johnstone, S. E., et al., Core transcriptional regulatory circuitry in human embryonic stem cells. Cell 2005, 122, 947–956.

[22] Lee, T. I., Jenner, R. G., Boyer, L. A., Guenther, M. G., et al., Control of developmental regulators by Polycomb in human embryonic stem cells. Cell 2006, 125, 301–313.

[23] Chen, X., Xu, H., Yuan, P., Fang, F., et al., Integration of external signaling pathways with the core transcriptional network in embryonic stem cells. Cell 2008, 133, 1106–1117.

[24] Moussaieff, A., Rouleau, M., Kitsberg, D., Cohen, M., et al., Glycolysis-mediated changes in acetyl-CoA and histone acetylation control the early differentiation of embryonic stem cells. Cell Metab. 2015, 21, 392–402.

[25] Ying, Q.-L., Wray, J., Nichols, J., Batlle-Morera, L., et al., The ground state of embryonic stem cell self-renewal. Nature 2008, 453, 519–523.

[26] Boroviak, T., Loos, R., Bertone, P., Smith, A., Nichols, J., The ability of inner-cell-mass cells to self-renew as embryonic stem cells is acquired following epiblast specification. Nat Cell Biol 2014, 16, 516–528.

[27] MacArthur, B. D., Sevilla, A., Lenz, M., Müller, F.-J., et al., Nanog-dependent feedback loops regulate murine embryonic stem cell heterogeneity. Nat Cell Biol 2012, 14, 1139–1147.

[28] Nichols, J., Zevnik, B., Anastassiadis, K., Niwa, H., et al., Formation of pluripotent stem cells in the mammalian embryo depends on the POU transcription factor Oct4. Cell 1998, 95, 379–391.

[29] Nakagawa, M., Koyanagi, M., Tanabe, K., Takahashi, K., et al., Generation of induced pluripotent stem cells without Myc from mouse and human fibroblasts. Nat Biotechnol 2008, 26, 101–106.

[30] Kim, J. B., Greber, B., Araúzo-Bravo, M. J., Meyer, J., et al., Direct reprogramming of human neural stem cells by OCT4. Nature 2009, 461, 649–643.

[31] Wang, J., Rao, S., Chu, J., Shen, X., et al., A protein interaction network for pluripotency of embryonic stem cells. Nature 2006, 444, 364–368.

[32] Liang, J., Wan, M., Zhang, Y., Gu, P., et al., Nanog and Oct4 associate with unique transcriptional repression complexes in embryonic stem cells. Nat Cell Biol 2008, 10, 731–739.

[33] Kim, J., Woo, A. J., Chu, J., Snow, J. W., et al., A Myc network accounts for similarities between embryonic stem and cancer cell transcription programs. Cell 2010, 143, 313–324.

[34] van den Berg, D.L.C., Snoek, T., Mullin, N. P., Yates, A., et al., An Oct4-centered protein interaction network in embryonic stem cells. Cell Stem Cell 2010, 6, 369–381.

[35] Pardo, M., Lang, B., Yu, L., Prosser, H., et al., An expanded Oct4 interaction network: implications for stem cell biology, development, and disease. Cell Stem Cell 2010, 6, 382–395.

[36] Gao, Z., Cox, J. L., Gilmore, J. M., Ormsbee, B. D., et al., Determination of protein interactome of transcription factor Sox2 in embryonic stem cells engineered for inducible expression of four reprogramming factors. J Biol Chem 2012, 287, 11384–11397.

[37] Loh, Y.-H., Wu, Q., Chew, J.-L., Vega, V. B., et al., The Oct4 and Nanog transcription network regulates pluripotency in mouse embryonic stem cells. Nat Genet 2006, 38, 431–440.

[38] Kim, J., Chu, J., Shen, X., Wang, J., Orkin, S. H., An extended transcriptional network for pluripotency of embryonic stem cells. Cell 2008, 132, 1049–1061.

[39] Ding, J., Xu, H., Faiola, F., Ma'ayan, A., Wang, J., Oct4 links multiple epigenetic pathways to the pluripotency network. Cell Res. 2012, 22, 155–167.

[40] Fidalgo, M., Faiola, F., Pereira, C.-F., Ding, J., et al., Zfp281 mediates Nanog autorepression through recruitment of the NuRD complex and inhibits somatic cell reprogramming. PNAS 2012, 109, 16202–16207.

[41] Efroni, S., Duttagupta, R., Cheng, J., Dehghani, H., et al., Global transcription in pluripotent embryonic stem cells. Cell Stem Cell 2008, 2, 437–447.

[42] Bernstein, B. E., Mikkelsen, T. S., Xie, X., Kamal, M., et al., A bivalent chromatin structure marks key developmental genes in embryonic stem cells. Cell 2006, 125, 315–326.

[43] Mikkelsen, T. S., Ku, M., Jaffe, D. B., Issac, B., et al., Genome-wide maps of chromatin state in pluripotent and lineage-committed cells. Nature 2007, 448, 553–560.

[44] Meissner, A., Mikkelsen, T. S., Gu, H., Wernig, M., et al., Genome-scale DNA methylation maps of pluripotent and differentiated cells. Nature 2008, 454, 766–770.

[45] Wang, Y., Medvid, R., Melton, C., Jaenisch, R., Blelloch, R., DGCR8 is essential for microRNA biogenesis and silencing of embryonic stem cell self-renewal. Nat Genet 2007, 39, 380–385.

[46] Judson, R. L., Babiarz, J. E., Venere, M., Blelloch, R., Embryonic stem cell-specific microRNAs promote induced pluripotency. Nat Biotechnol 2009, 27, 459–461.

[47] Xu, N., Papagiannakopoulos, T., Pan, G., Thomson, J. A., Kosik, K. S., MicroRNA-145 regulates OCT4, SOX2, and KLF4 and represses pluripotency in human embryonic stem cells. Cell 2009, 137, 647–658.

[48] Leonardo, T. R., Schultheisz, H. L., Loring, J. F., Laurent, L. C., The functions of microRNAs in pluripotency and reprogramming. Nat Cell Biol 2012, 14, 1114–1121.

[49] Ingolia, N. T., Lareau, L. F., Weissman, J. S., Ribosome profiling of mouse embryonic stem cells reveals the complexity and dynamics of mammalian proteomes. Cell 2011, 147, 789–802.

[50] Yue, F., Cheng, Y., Breschi, A., Vierstra, J., et al., A comparative encyclopedia of DNA elements in the mouse genome. Nature 2014, 515, 355–364.

[51] Williams, R. L., Hilton, D. J., Pease, S., Willson, T. A., et al., Myeloid leukaemia inhibitory factor maintains the developmental potential of embryonic stem cells. Nature 1988, 336, 684–687.

[52] Smith, A. G., Heath, J. K., Donaldson, D. D., Wong, G. G., et al., Inhibition of pluripotential embryonic stem cell differentiation by purified polypeptides. Nature 1988, 336, 688–690.

[53] Ying, Q.-L., Nichols, J., Chambers, I., Smith, A., BMP induction of Id proteins suppresses differentiation and sustains embryonic stem cell self-renewal in collaboration with STAT3. Cell 2003, 115, 281–292.

[54] Burdon, T., Stracey, C., Chambers, I., Nichols, J., Smith, A., Suppression of SHP-2 and ERK signalling promotes self-renewal of mouse embryonic stem cells. Dev. Biol. 1999, 210, 30–43.

[55] Stavridis, M. P., Lunn, J. S., Collins, B. J., Storey, K. G., A discrete period of FGF-induced Erk1/2 signalling is required for vertebrate neural specification. Development 2007, 134, 2889–2894.

[56] Kunath, T., Saba-El-Leil, M.K., Almousailleakh, M., Wray, J., et al., FGF stimulation of the Erk1/2 signalling cascade triggers transition of pluripotent embryonic stem cells from self-renewal to lineage commitment. Development 2007, 134, 2895–2902.

[57] Amit, M., Carpenter, M. K., Inokuma, M. S., Chiu, C. P., et al., Clonally derived human embryonic stem cell lines maintain pluripotency and proliferative potential for prolonged periods of culture. Dev. Biol. 2000, 227, 271–278.

[58] Xu, R.-H., Peck, R. M., Li, D. S., Feng, X., et al., Basic FGF and suppression of BMP signaling sustain undifferentiated proliferation of human ES cells. Nat Methods 2005, 2, 185–190.

[59] Sato, N., Meijer, L., Skaltsounis, L., Greengard, P., Brivanlou, A. H., Maintenance of pluripotency in human and mouse embryonic stem cells through activation of Wnt signaling by a pharmacological GSK-3-specific inhibitor. Nat Med 2004, 10, 55–63.

[60] Gruber, A. J., Grandy, W. A., Balwierz, P. J., Dimitrova, Y. A., et al., Embryonic stem cell-specific microRNAs contribute to pluripotency by inhibiting regulators of multiple differentiation pathways. Nucleic Acids Res 2014, 42, 9313–9326.

[61] Pelkmans, L., Using cell-to-cell variability-a new era in molecular biology. Science 2012, 336, 425–426.

[62] Bendall, S. C., Nolan, G. P., Roederer, M., Chattopadhyay, P. K., A deep profiler's guide to cytometry. Trends Immunol. 2012, 33, 323–332.

[63] Bandura, D. R., Baranov, V. I., Ornatsky, O. I., Antonov, A., et al., Mass cytometry: technique for real time single cell multitarget immunoassay based on inductively coupled plasma time-of-flight mass spectrometry. Anal. Chem. 2009, 81, 6813–6822.

[64] Di Palma, S., Bodenmiller, B., Unraveling cell populations in tumors by single-cell mass cytometry. Curr Opin Biotechnol 2015, 31, 122–129.

[65] Femino, A. M., Fay, F. S., Fogarty, K., Singer, R. H., Visualization of single RNA transcripts in situ. Science 1998, 280, 585–590.

[66] Raj, A., van den Bogaard, P., Rifkin, S. A., van Oudenaarden, A., Tyagi, S., Imaging individual mRNA molecules using multiple singly labeled probes. Nat Methods 2008, 5, 877–879.

[67] White, A. K., Vaninsberghe, M., Petriv, O. I., Hamidi, M., et al., High-throughput microfluidic single-cell RT-qPCR. PNAS 2011, 108, 13999–14004.

[68] Tang, F., Barbacioru, C., Wang, Y., Nordman, E., et al., mRNA-Seq whole-transcriptome analysis of a single cell. Nat Methods 2009, 6, 377–382.

[69] Hoppe, P. S., Coutu, D. L., Schroeder, T., Single-cell technologies sharpen up mammalian stem cell research. Nat Cell Biol 2014, 16, 919–927.

[70] Sachs, K., Perez, O., Pe'er, D., Lauffenburger, D. A., Nolan, G. P., Causal protein-signaling networks derived from multiparameter single-cell data. Science 2005, 308, 523–529.

[71] Chambers, I., Silva, J., Colby, D., Nichols, J., et al., Nanog safeguards pluripotency and mediates germline development. Nature 2007, 450, 1230–1234.

[72] Toyooka, Y., Shimosato, D., Murakami, K., Takahashi, K., Niwa, H., Identification and characterization of subpopulations in undifferentiated ES cell culture. Development 2008, 135, 909–918.

[73] Hayashi, K., Lopes, S.M.C. de S., Tang, F., Surani, M. A., Dynamic equilibrium and heterogeneity of mouse pluripotent stem cells with distinct functional and epigenetic states. Cell Stem Cell 2008, 3, 391–401.

[74] Faddah, D. A., Wang, H., Cheng, A. W., Katz, Y., et al., Single-cell analysis reveals that expression of nanog is biallelic and equally variable as that of other pluripotency factors in mouse ESCs. Cell Stem Cell 2013, 13, 23–29.

[75] Kolodziejczyk, A. A., Kim, J. K., Tsang, J.C.H., Ilicic, T., et al., Single Cell RNA-Sequencing of Pluripotent States Unlocks Modular Transcriptional Variation. Cell Stem Cell 2015, 17, 471–485.

[76] Thattai, M., van Oudenaarden, A., Intrinsic noise in gene regulatory networks. PNAS 2001, 98, 8614–8619.

[78] Becskei, A., Kaufmann, B. B., van Oudenaarden, A., Contributions of low molecule number and chromosomal positioning to stochastic gene expression. Nat Genet 2005, 37, 937–944.

[78] Trott, J., Hayashi, K., Surani, A., Babu, M. M., Martinez Arias, A., Dissecting ensemble networks in ES cell populations reveals micro-heterogeneity underlying pluripotency. Mol Biosyst 2012, 8, 744–752.

[79] Xu, H., Ang, Y.-S., Sevilla, A., Lemischka, I. R., Ma'ayan, A., Construction and validation of a regulatory network for pluripotency and self-renewal of mouse embryonic stem cells. PLoS Comput Biol 2014, 10, e1003777.

[80] MacArthur, B. D., Collective dynamics of stem cell populations. PNAS 2014, 111, 3653–3654.

[81] Luscombe, N. M., Babu, M. M., Yu, H., Snyder, M., et al., Genomic analysis of regulatory network dynamics reveals large topological changes. Nature 2004, 431, 308–312.

[82] Davidson, E. H., Levine, M. S., Properties of developmental gene regulatory networks. PNAS 2008, 105, 20063–20066.

[83] Müller, F.-J., Laurent, L. C., Kostka, D., Ulitsky, I., et al., Regulatory networks define phenotypic classes of human stem cell lines. Nature 2008, 455, 401–405.

[84] Cahan, P., Li, H., Morris, S. A., Lummertz da Rocha, E., et al., CellNet: network biology applied to stem cell engineering. Cell 2014, 158, 903–915.

[85] Chazaud, C., Yamanaka, Y., Pawson, T., Rossant, J., Early lineage segregation between epiblast and primitive endoderm in mouse blastocysts through the Grb2-MAPK pathway. Dev Cell 2006, 10, 615–624.

[86] Yamanaka, Y., Lanner, F., Rossant, J., FGF signal-dependent segregation of primitive endoderm and epiblast in the mouse blastocyst. Development 2010, 137, 715–724.

[87] Plusa, B., Piliszek, A., Frankenberg, S., Artus, J., Hadjantonakis, A.-K., Distinct sequential cell behaviours direct primitive endoderm formation in the mouse blastocyst. Development 2008, 135, 3081–3091.

[88] Meilhac, S. M., Adams, R. J., Morris, S. A., Danckaert, A., et al., Active cell movements coupled to positional induction are involved in lineage segregation in the mouse blastocyst. Dev. Biol. 2009, 331, 210–221.

[89] Yamanaka, Y., Ralston, A., Stephenson, R. O., Rossant, J., Cell and molecular regulation of the mouse blastocyst. Dev Dyn 2006, 235, 2301–2314.

[90] Morris, S. A., Teo, R.T.Y., Li, H., Robson, P., et al., Origin and formation of the first two distinct cell types of the inner cell mass in the mouse embryo. PNAS 2010, 107, 6364–6369.

[91] Morris, S. A., Graham, S.J.L., Jedrusik, A., Zernicka-Goetz, M., The differential response to Fgf signalling in cells internalized at different times influences lineage segregation in preimplantation mouse embryos. Open Biol 2013, 3, 130–104.

[92] Song, J., Saha, S., Gokulrangan, G., Tesar, P. J., Ewing, R. M., DNA and chromatin modification networks distinguish stem cell pluripotent ground states. Mol. Cell Proteomics 2012, 11, 1036–1047.

[93] Guo, G., Yang, J., Nichols, J., Hall, J. S., et al., Klf4 reverts developmentally programmed restriction of ground state pluripotency. Development 2009, 136, 1063–1069.

[94] Takashima, Y., Guo, G., Loos, R., Nichols, J., et al., Resetting transcription factor control circuitry toward ground-state pluripotency in human. Cell 2014, 158, 1254–1269.

[95] Rossant, J., Tam, P.P.L., Blastocyst lineage formation, early embryonic asymmetries and axis patterning in the mouse. Development 2009, 136, 701–713.

[96] Habib, S. J., Chen, B.-C., Tsai, F.-C., Anastassiadis, K., et al., A localized Wnt signal orients asymmetric stem cell division in vitro. Science 2013, 339, 1445–1448.

[97] Tsankov, A. M., Gu, H., Akopian, V., Ziller, M. J., et al., Transcription factor binding dynamics during human ES cell differentiation. Nature 2015, 518, 344–349.

[98] Dixon, J. R., Jung, I., Selvaraj, S., Shen, Y., et al., Chromatin architecture reorganization during stem cell differentiation. Nature 2015, 518, 331-–336.

[99] Apostolou, E., Hochedlinger, K., Chromatin dynamics during cellular reprogramming. Nature 2013, 502, 462–471.

[100] Heng, J.-C.D., Feng, B., Han, J., Jiang, J., et al., The nuclear receptor Nr5a2 can replace Oct4 in the reprogramming of murine somatic cells to pluripotent cells. Cell Stem Cell 2010, 6, 167–174.

[101] Li, R., Liang, J., Ni, S., Zhou, T., et al., A mesenchymal-to-epithelial transition initiates and is required for the nuclear reprogramming of mouse fibroblasts. Cell Stem Cell 2010, 7, 51–63.

[102] Stadtfeld, M., Maherali, N., Breault, D. T., Hochedlinger, K., Defining molecular cornerstones during fibroblast to iPS cell reprogramming in mouse. Cell Stem Cell 2008, 2, 230–240.

[103] Mikkelsen, T. S., Hanna, J., Zhang, X., Ku, M., et al., Dissecting direct reprogramming through integrative genomic analysis. Nature 2008, 454, 49–55.

[104] Wernig, M., Lengner, C. J., Hanna, J., Lodato, M. A., et al., A drug-inducible transgenic system for direct reprogramming of multiple somatic cell types. Nat Biotechnol 2008, 26, 916–924.

[105] Hanna, J., Saha, K., Pando, B., van Zon, J., et al., Direct cell reprogramming is a stochastic process amenable to acceleration. Nature 2009, 462, 595–601.

[106] MacArthur, B. D., Please, C. P., Oreffo, R.O.C., Stochasticity and the molecular mechanisms of induced pluripotency. PLoS ONE 2008, 3, e3086.

[107] Yamanaka, S., Elite and stochastic models for induced pluripotent stem cell generation. Nature 2009, 460, 49–52.

[108] Rais, Y., Zviran, A., Geula, S., Gafni, O., et al., Deterministic direct reprogramming of somatic cells to pluripotency. Nature 2013, 502, 65–70.

[109] Buganim, Y., Faddah, D. A., Cheng, A. W., Itskovich, E., et al., Single-cell expression analyses during cellular reprogramming reveal an early stochastic and a late hierarchic phase. Cell 2012, 150, 1209–1222.

[110] Zunder, E. R., Lujan, E., Goltsev, Y., Wernig, M., Nolan, G. P., A continuous molecular roadmap to iPSC reprogramming through progression analysis of single-cell mass cytometry. Cell Stem Cell 2015, 16, 323–337.

[111] Lujan, E., Zunder, E. R., Ng, Y. H., Goronzy, I. N., et al., Early reprogramming regulators identified by prospective isolation and mass cytometry. Nature 2015.

[112] Ivanova, N., Dobrin, R., Lu, R., Kotenko, I., et al., Dissecting self-renewal in stem cells with RNA interference. Nature 2006, 442, 533–538.

[113] Lu, R., Markowetz, F., Unwin, R. D., Leek, J. T., et al., Systems-level dynamic analyses of fate change in murine embryonic stem cells. Nature 2009, 462, 358–362.

[114] Wang, Z., Oron, E., Nelson, B., Razis, S., Ivanova, N., Distinct Lineage Specification Roles for NANOG, OCT4, and SOX2 in Human Embryonic Stem Cells. Stem Cell 2012, 10, 440–454.

[115] Kalman, R. E., Mathematical description of linear dynamical systems. Journal of the Society for Industrial & Applied … 1963, 1, 152–192.

[116] Liu, Y.-Y., Slotine, J.-J., Barabási, A.-Á., Controllability of complex networks. Nature 2011, 473, 167–173.

[117] Scutari, M., Learning Bayesian Networks with the bnlearn R Package. JStat Softw 2010, 35, 1–22.

[118] Lèbre, S., Becq, J., Devaux, F., Stumpf, M. P., Lelandais, G., Statistical inference of the time-varying structure of gene-regulation networks. BMC Syst Biol 2010, 4, 130–16.

[119] Singer, Z. S., Yong, J., Tischler, J., Hackett, J. A., et al., Dynamic heterogeneity and DNA methylation in embryonic stem cells. 2014, 55, 319–331.

[120] Streets, A. M., Zhang, X., Cao, C., Pang, Y., et al., Microfluidic single-cell whole-transcriptome sequencing. PNAS 2014, 111, 7048–7053.

[121] Klein, A.M., Mazutis, L., Akartuna, I., Tallapragada, N., et al., Droplet barcoding for single-cell transcriptomics applied to embryonic stem cells. Cell 2015, 161, 1187–1201.

